# Remote targeted electrical stimulation

**DOI:** 10.1101/2021.10.09.463785

**Authors:** Rahul Cheeniyil, Jan Kubanek

## Abstract

The ability to generate electric fields in deep tissues remotely, without surgically implanting electrodes, could transform diagnoses and treatments of nervous system disorders. Here, we show that focal electrostimulation effects can be elicited remotely by combining two noninvasive forms of energies—magnetic and focused ultrasonic fields. The approach, based in the Lorentz equation and referred to as Lstim, electrically stimulates specified tissue targets with the precision of deep brain or spinal cord stimulation, but does not require electrode implantation. Lstim potentiated the responses of human nerves, enhancing the neuromodulatory effects of ultrasound by 74% on average. The effects showed a double dissociation—a significant and substantial increase in nociceptive responses, yet a significant reduction in tactile responses. In line with the Lorentz equation, Lstim was only observed when nerves were oriented perpendicularly to the magnetic and ultrasonic fields. A sham condition showed no effects. Both the ultrasonic and the induced electric fields were well below the respective safety indices, and no detrimental effects were detected. Lstim uniquely integrates noninvasiveness, sharp focus, and the efficacy of electrical stimulation. The approach has the potential to provide a noninvasive tool to dissect brain function in humans and to diagnose the neural circuits involved in nervous system disorders. Moreover, this effect should be taken into account when ultrasound is applied inside MRI.

## Introduction

Nearly one in four people lives with a significant neurological or psychiatric disorder (Lancet, 2017; Ahrnsbrak et al., 2017). Approximately one in three patients across neurological and psychiatric conditions does not respond to drugs or has intolerable side effects (Bystritsky, 2006; Al-Harbi, 2012; Jaffe et al., 2019; Lyons and Pahwa, 2004; Zesiewicz et al., 2010; Elias et al., 2013; Elias et al., 2016; Ferguson, 2001; Karceski, 2007; Louis et al., 2010). Bioelectronic medicine, also referred to as neuromodulation, provides these patients with new treatment options, promising to treat nervous system disorders at their neural source (Perlmutter and Mink, 2006; Arle and Shils, 2017).

Surgery-based electrostimulation approaches, including deep brain stimulation, spinal cord stimulation, sacral nerve stimulation, and vagal nerve stimulation have provided relief to many patients with specific conditions, including movement disorders (Larson, 2014), chronic pain (Lempka and Patil, 2018), bladder control (van Balken et al., 2004), and epilepsy (Elliott et al., 2011), respectively. However, these surgical approaches are costly and incur risks and side effects (Tonge et al., 2015; Bergey et al., 2015; Shah et al., 2015; Sinai et al., 2019; Giordano et al., 2020). Consequently, these approaches are generally limited to an established site of implantation.

Noninvasive approaches, which have rested on electrical, magnetic, electromagnetic, and ultrasonic fields, have much greater flexibility in that they do not incur additional risk to subjects when modulating multiple sites. However, these approaches do not currently have the necessary spatial resolution to modulate specific neural circuits at depth.

Electric fields generated with current noninvasive approaches, including electroconvulsive therapy (ECT) or transcranial direct or alternating current stimulation (tDCS, tACS) are relatively broad (Lisanby, 2007; CAUMO et al., 2012; Herrmann et al., 2013). Consequently, the sizable activation of the brain associated with ECT often results in cognitive side effects (Ingram et al., 2008). The spatial resolution of these methods can be improved using spatially interfering fields (Nemec, 1959; Grossman et al., 2017), but the resulting fields are still broad within respect to the dimensions of neural circuits.

Transcranial magnetic stimulation (TMS) uses pulses of magnetic fields to noninvasively induce electric fields in the brain. TMS can produce appreciable effects in the cortex and ameliorate symptoms of depression (George et al., 2000), but the approach cannot directly and focally modulate deep brain regions.

Electromagnetic waves currently cannot be used to modulate deep brain targets in a focal manner. At high frequencies (light or infrared), electromagnetic waves are severely attenuated by the skull or superficial tissue layers (McCormick et al., 1992). At lower frequencies, the waves (microwaves) can penetrate into depth, but microwaves at the relevant neuromodulatory doses damage mitochondria and possibly other cellular structures (McRee and Wachtel, 1980; Hao et al., 2015). At yet lower frequencies (radio range), the wavelength is too broad—dozens of centimeters or meters—to allow for focal stimulation (Lustenberger et al., 2013).

Low-intensity focused ultrasound combines depth penetration and safe application. Ultrasound can effectively modulate excitable cells at high frequencies—above 10 MHz—at which there are strong radiation forces that mechanically displace membranes and activate ion channels (Menz et al., 2013; Kubanek et al., 2016; Kubanek et al., 2018; Prieto et al., 2018). However, ultrasound at such high frequencies is severely attenuated by the human skull (Fry, 1977; Fry and Barger, 1978); for this reason, frequencies below 1 MHz have been used for transcranial therapies (Kubanek, 2018). Ultrasound can modulate excitable structures also at lower frequencies (Naor et al., 2016; Blackmore et al., 2019), but strong and reproducible effects, under safe ultrasound levels, remain elusive.

To fill in the current technological gap, we developed and validated an approach that combines the noninvasiveness and targeting capabilities of low-intensity ultrasound with the well-understood and potent effects of electrical stimulation (Cohen and Newsome, 2004; Perlmutter and Mink, 2006; Penfield and Boldrey, 1937; Fried et al., 1991; Suthana et al., 2012). The approach combines two forms of noninvasive energies—ultrasonic and magnetic fields (Figure 1). The idea is based on the fact that motion of a charged molecule *q* in magnetic field 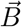 generates Lorentz force 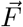 on the molecule: 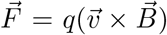. The generated electrical field has intensity 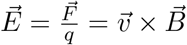. To achieve localized electric field, the movement 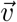 should occur only in the target of interest. The critical insight is that this can be achieved using focused ultrasonic waves. In particular, focused ultrasound induces in its target pressure *P*. This pressure leads to a displacement of molecules at the target, with velocity 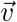 that points in the direction of the propagating wave. In biological tissues, the speed of the displacement is proportional to the pressure at target, 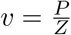, where *Z* is a tissue constant, “acoustic impedance”. Thus, acoustic waves delivered into a target perpendicularly to a magnetic field produce in the target electric field intensity 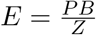. This intensity points in the direction that is perpendicular to both constituents. As a consequence of this electric field, positively and negatively charged molecules are pulled in opposite directions, inducing electric currents. (Ultrasonic fields alone displace positively and negatively charged molecules in the same direction. Therefore, no gradient of charge and so no electric field would be created. The magnetic field is a critical addition.) We refer to this stimulation as “Lstim” given its origin in the Lorentz equation and its electrical and local nature.

**Figure 1.**
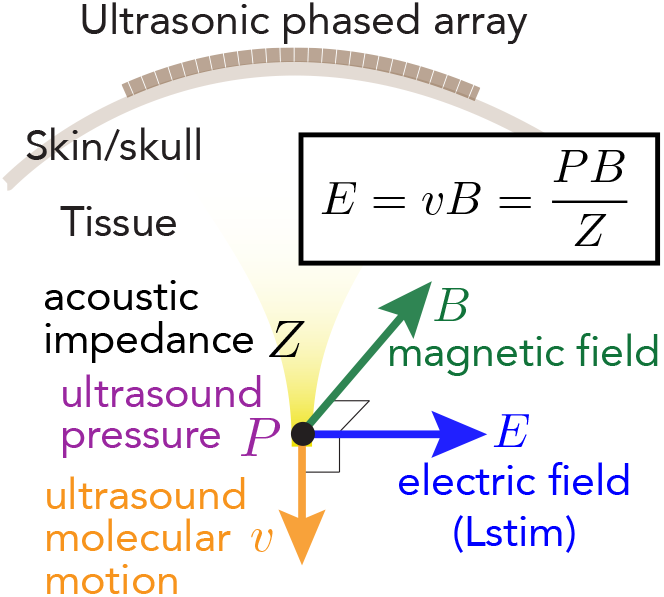
Electrical stimulation without electrodes. An MRI magnet generates magnetic field *B*. An ultrasonic transducer array programmatically focuses ultrasound into a neural circuit of interest. An ultrasound wave, focused into a target with acoustic impedance *Z*, induces in the target motions of molecules with velocity 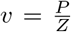. The pressure *P* (and so the velocity *v*) are maximal at the target. When the wave is emitted in a direction perpendicular to *B*, so that the velocity vector is perpendicular to *B*, the target experiences localized electric field 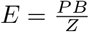.

A proof of concept of Lstim has been demonstrated in a study that applied weak ultrasound pulses (≤ 83 kPa) to mouse motor cortex in a weak magnetic field (0.15 T). There were small but significant changes in evoked motor responses of the animals (Wang et al., 2019). Given an effect at a very weak electric field (about 8.3 mV/m (Wang et al., 2019)), we predict that Lstim using a strong magnetic field will (7 T) will have substantial effects on neural structures.

Mechanistically, the generated electric field 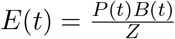 has, for static magnetic field *B*, the same frequency as that of the applied ultrasound, which is generally in the hundreds of kilohertz range. Stimulation at high frequencies rests on a “Gildermeister effect”, or integration of membrane potential toward a threshold (Ward, 2009). Moreover, high-frequency electrical stimuli produce a reliable “onset response”—a transient neural response following stimulus onset (Ward, 2009; Grossman et al., 2017). Therefore, when a stimulus is pulsed, we can expect that the onset response will occur within each pulse, thus maximizing the response.

## Results

We applied Lstim to the peripheral nervous system of 18 human subjects (Figure 2). The stimulation was performed inside and outside a 7 T scanner, with order randomized across the subjects. We tested a total of 9 distinct stimuli—3 distinct amplitudes and 3 distinct waveforms, and included a sham condition (Methods). The ultrasound was delivered into the tissues through a water-filled coupling cone (Figure 2a).

**Figure 2.**
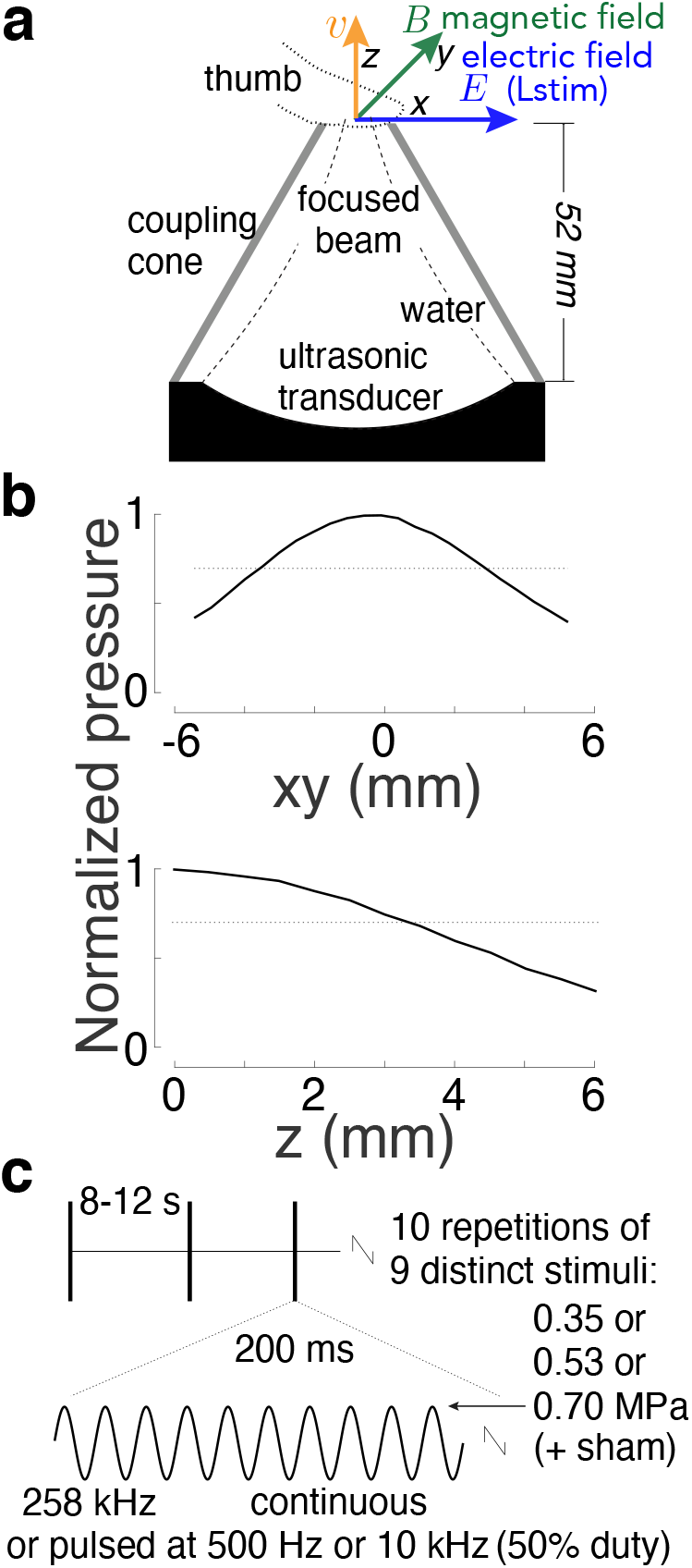
Apparatus and stimuli. a) Apparatus. A focused ultrasound transducer delivered a 258 kHz stimulus into a subject’s thumb using a coupling cone filled with degassed water. The stimulation was performed inside a 7 T scanner or 3 m away from it. Subjects were instructed to place the thumb in the direction perpendicular to the magnetic and ultrasonic fields, to maximize Lstim-based effects. b) Peaknormalized ultrasound pressure field. The pressure profile was averaged over the x and y dimensions. The dotted lines show the 0.707 (0.5) pressure (intensity) levels to characterize the fields using full-width-at-half-maximum values. The full-width-at-half-maximum (FWHM) diameter was 6.5 mm, and focal length 3.3 mm. c) Stimuli. Each subject experienced 10 repetitions of 10 distinct stimuli, including sham. The stimuli, 200 ms in duration, were selected randomly and delivered each 8-12 seconds. We tested 3 pressure levels and continuous and pulsed (500 Hz, 10 kHz frequency, 50% duty) stimuli.

The individual stimuli (Figure 2c) were delivered randomly every 8-12 seconds. The subjects were asked to report any nociceptive or tactile sensation. A nociceptive sensation results from an activation of free nerve endings in the skin (Dubin et al., 2010) and so constitutes a metric of neural activation. The ultrasonic stimuli were safely within the 510(k) FDA safety indices (FDA, 2019), and the induced electric fields were safely below the recommended charge density (Cogan et al., 2016) of 30 *μ*C/cm^2^ (Figure 3).

**Figure 3.**
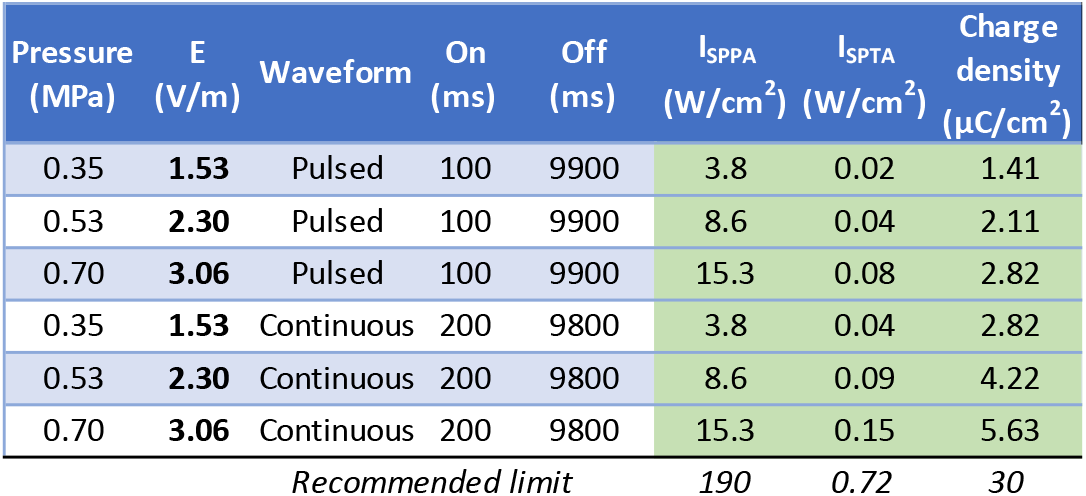
Stimuli and their safety. The study used 9 distinct stimuli: 3 levels of pressure, and 3 distinct waveforms, one continuous (100% duty) and two pulsed (50% duty). All stimuli were 200 ms in duration and were delivered every 10 s on average. The study followed the FDA 510(k) Track 3 recommendations (FDA, 2019): peak intensity *I*_SPPA_ and time-average intensity *I*_SPTA_. *E* is the induced peak Lstim intensity in a 7 T magnetic field. The computation of the charge density assumes brain conductivity of 0.26 S/m (Koessler et al., 2017). Electrical stimulation should ideally not exceed charge density of 30 *μ*C/cm^2^ (Cogan et al., 2016). Green entries indicate that all stimuli are well within the recommended levels.

We found that ultrasound delivered into the nerves of the subjects’ finger inside the magnetic field substantially enhanced the magnitude of nociceptive responses (Figure 4). Across all pressure levels and waveforms, Lstim increased the magnitude of nociceptive responses by 74%. There was a double dissociation of the effects with respect to magnetic field and sensation kind (two-way ANOVA, magnetic field × sensation interaction, *p* < 0.001; *F*(1, 644) = 13.20). The effect was similar when the responses were not scaled by their intensity (*p* < 0.001; *F* (1, 644) = 13.93). Pairwise tests showed that the increase in the nociceptive responses (*p* = 0.0059; *t*(17) = 3.14, paired two-sided t-test) as well as the decrease in tactile responses (*p* = 0.0033; *t*(17) = −3.41) were significant. The effects were similar when the responses were not scaled by their intensity (*p* = 0.0037; *t*(17) = 3.36 and *p* = 0.0029; *t*(17) = −3.47, respectively).

**Figure 4.**
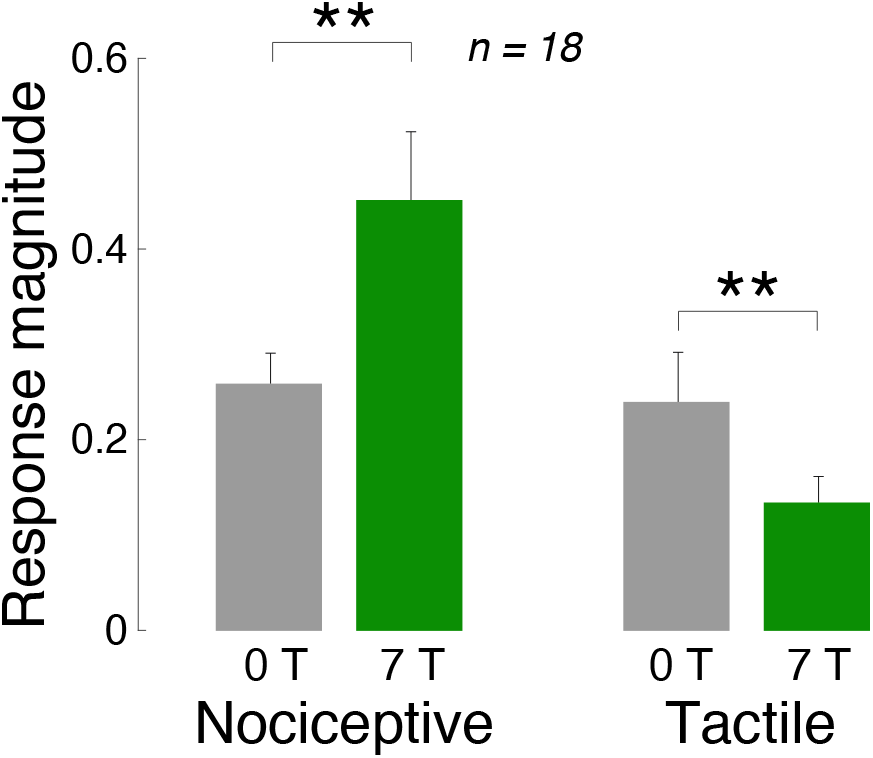
Magnetic field combined with focused ultrasound remotely produces targeted neurostimulatory effects. Mean s.e.m. response magnitude (see Methods) for ultrasound alone (0 T) and ultrasound combined with magnetic field (7 T), separately for nociceptive (left) and tactile (right) responses. Data were pooled over all stimuli tested. The double stars indicate effects significant at *p* < 0.01.

We next specifically analyzed the nociceptive responses, which reflect an activation of nerves or nerve endings (Dubin et al., 2010). Figure 5 shows the dependence of all stimuli on the presence or absence of magnetic field, separately for each pressure. The figure confirms the findings of Figure 4 that the magnetic field amplifies the nociceptive responses. We assessed the effects using a full, three-way ANOVA model with factors magnetic field, ultrasound pressure, stimulus waveform, and all possible interactions (Table 1). Indeed, the effect of magnetic field was significant also in this analysis (*p* < 0.001, *F* (1, 408) = 18.55).

**Table 1.** Summary of the effects. The effects of magnetic field (*M*), ultra-sound pressure (*P*), and stimulus waveform (*W*; continuous or pulsed) on the frequency of nociceptive (left column) and tactile responses (right column). These effects were assessed using a three-way ANOVA that featured the three main effects and all possible interactions. Bold entries are significant (*p* < 0.05).

|  | Nociceptive | Tactile |
| --- | --- | --- |
| $M$ | **< 0.001** | **0.0042** |
| $P$ | **< 0.001** | **< 0.001** |
| $W$ | **< 0.001** | **< 0.001** |
| $M \times P$ | **0.0012** | 0.23 |
| $M \times W$ | **0.043** | **0.040** |
| $P \times W$ | **< 0.001** | **< 0.001** |
| $M \times P \times W$ | 0.54 | 0.83 |

**Figure 5.**
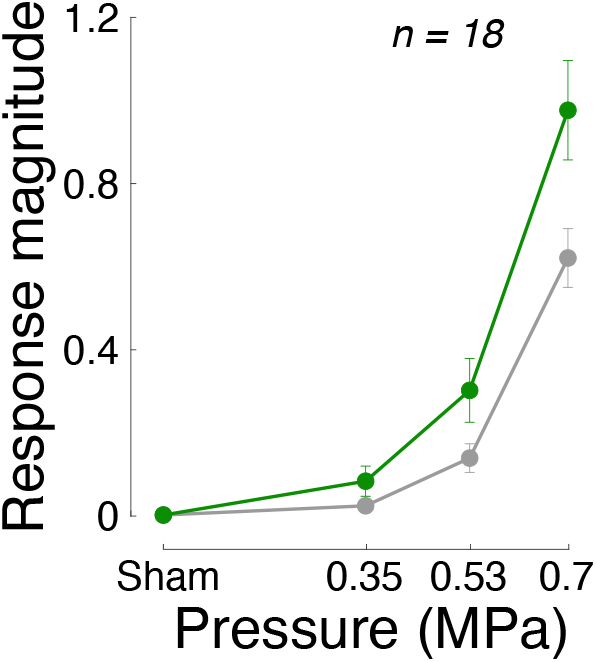
The stimulatory effects as a function of ultrasound pressure. Mean ± s.e.m. magnitude of nociceptive responses as a function of magnetic field and the pressure of ultrasound at target. Data were pooled over all waveforms.

Lstim produces focused electric fields at ultrasound targets according to 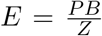. In this equation, the effect increases with the ultrasound pressure *P*. Therefore, the higher the ultrasound pressure, the stronger the induced electric fields, and the stronger the nociceptive responses we should observe, in addition to any neuromodulatory effects of ultrasound alone. In line with this expectation, we found a significant interaction between the magnetic field and ultrasound pressure (*p* = 0.0012, *F* (3, 408) = 5.41).

We summarize the effects of all factors and interactions in Table 1. With respect to nerve activation, as assessed by the nociceptive responses, there was a significant interaction between magnetic field and the stimulus waveform (*p* = 0.043, *F*(2, 408) = 3.16). As expected (see Introduction), the contrast between Lstim and ultrasound only was higher when the ultrasound was pulsed. Specifically, averaged across all pressures, the response frequency ratio (7T versus 0T) for the continuous waveform was 1.61, compared to 1.85 and 3.85 for the pulsed 500 Hz and 10 kHz waveforms, respectively.

If the reported effects are indeed due to the induction of localized electric field, as governed by the Lorentz electromotive force equation, they should strongly depend on the orientation of the finger with respect to the electric field. Specifically, electric fields can effectively stimulate nerves if their gradients point along nerves, as opposed to across (Rattay, 1999). To test this, 4 subjects were asked to place their thumb on the aperture 1) perpendicularly to the magnetic field (up until now) 2) in parallel with the magnetic field. We found (Figure 6) that these conditions significantly modulated the responses (*p* = 0.041, *F* (2, 33) = 3.5). However, as expected, the effect was specific to the perpendicular geometry; there was no effect for the parallel geometry (*p* = 0.88, *t*(3) = 0.17, paired two-sided t-test).

**Figure 6.**
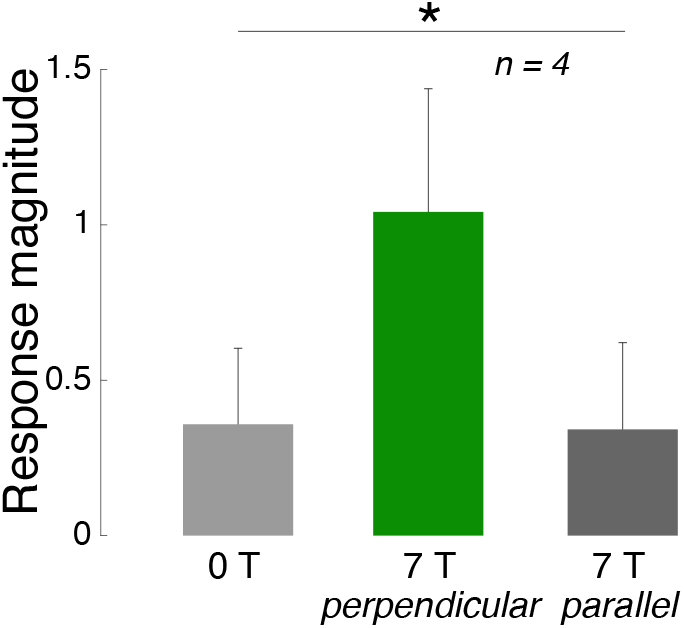
Lstim stimulates nerves in an orientation-specific manner. Mean ± s.e.m. magnitude of nociceptive responses as a function of the orientation of the induced electric field with respect to the subjects’ nerves. The neuromodulatory effects are maximized when nerves in the subjects’ thumb are aligned with the induced electric field (green). Data were pooled over all waveforms tested. Star: the modulation by the magnetic field and its orientation was significant (*p* < 0.05).

## Discussion

In this study, we applied and tested an approach that induces focal electric fields in remote targets. The approach, Lstim, rests on the Lorentz effect—movement of charged particles in strong magnetic field. Critically, in the Lstim embodiment of the Lorentz effect, the movement is induced by cycle-by-cycle displacement of particles at the target of focused ultrasound. We found that the combination of ultrasonic and magnetic fields safely produced effects that were about 1.7 times stronger compared to those induced by ultrasound alone. As expected by the governing equation (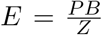), we found that the effects scaled as a function of the ultrasound pressure. In addition, the effects followed the direction-specific Lorentz equation. No effects were observed for the sham condition. Together, these findings confirm the hypothesis that a remote generation of focal electric fields by combining ultrasonic and magnetic fields is indeed feasible.

The produced electric field is deterministic, governed solely by the analytic expression 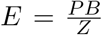 for cases in which the ultrasonic and magnetic fields are perpendicular. (Otherwise, the equation is multiplied by the sine of the angle between the two fields.) The acoustic pressure and its waveform *P* (*t*) can be measured for a given transducer hardware using a hydrophone. The static magnetic field of MRI scanners is near-homogeneous within an organ (e.g., brain), and so *B* is also a deterministic quantity (e.g., 7 T). The acoustic impedance tissue constant *Z* is close to water and varies only within about 10% across soft tissues (Cobbold, 2006; Azhari, 2010). For the brain, the acoustic impedance is about 1.6 MRayl (Azhari, 2010). Using this equation, we can therefore accurately compute the generated electric field intensity. For example, a 0.7 MPa stimulus at a target inside a 7 T MRI scanner, used here, evoked peak electric intensity of about 3.1 V/m. A field of 3 V/m is deemed to be high enough to appreciably influence neural activity (Francis et al., 2003; Liu et al., 2018). Fields as low 0.3 V/m have been shown to modulate neuronal spiking (Francis et al., 2003). For instance, transcranial electrical stimulation, for the generally accepted maximum current of 2 mA, produces about 0.28 V/m (95th percentile) in the brain (Huang et al., 2017). With Lstim, we elicited up to 11 times that intensity while complying with the applicable safety indices (Figure 3).

The effects on nerves were observed already for a relative low ultrasound pressure (0.7 MPa in amplitude at focus). We limited the highest pressure level to 0.7 MPa to avoid strong nociceptive sensations. No subject lifted their finger from the aperture during the experiment, and no harmful effects were detected by the experimenter or reported by the subjects. In regard to safety, the FDA 510(k) *I*_SPPA_ ultrasound index would allow us to apply up to 2.4 MPa (< 190 W/cm^2^ in soft tissues). At these levels, still considered safe, the effects of Lstim could be expected to be more than 3 times stronger than those reported here. Moreover, for the relatively low frequencies such as those used here, it is conceivable that ultrasound amplitudes higher than 2.4 MPa could be applied in brief pulses without a risk of heating or cavitation (Nightingale et al., 2015). If even stronger effects are needed for certain applications, the stimulation could be performed in stronger magnetic fields *B*. For example, in a 11 T scanner, a 2.4 MPa ultrasound stimulus would produce a peak intensity of about 16.5 V/m at focus, more than 5 times higher than that produced here. In addition, neurons are generally more sensitive to electrical stimuli than nerves. Therefore, Lstim is expected to elicit particularly strong effects when applied to neural circuits.

In fact, Lstim may have been adventitiously at play in the brain already. We revisited the ultrasonic neuromodulation literature (Naor et al., 2016; Kubanek, 2018; Blackmore et al., 2019) with a special focus on the presence of magnetic field. We now make the observation that when subjects are inside MRI, ultrasound can elicit strong effects including phosphenes (Lee et al., 2016) and can have appreciable effects on neural activity in humans (Ai et al., 2018). The unbeknown presence of Lstim may thus explain the notable effects in some “ultrasonic neuromodulation” studies.

In addition, several recent studies applied ultrasound of comparable frequency and pressure level to specific regions of the monkey brain inside a 3 T magnetic field. It has been found that only 40 seconds of such stimulation could dissociate network connectivity in deep brain regions including the amygdala, and these neuroplastic effects outlive the stimulation for about two hours (Verhagen et al., 2019; Fouragnan et al., 2019; Khalighinejad et al., 2020). In many of the cases, the ultrasound was applied to the brain near-perpendicularly to the magnetic field. Given the findings of the present study, therefore, these studies have, at least in part, invoked Lstim. To what extent Lstim contributed to the effects should be investigated in future studies.

There is additional evidence that the effects of Lstim can be durable and so potentially also useful for treatments. Specifically, when subjects move their head near an MRI scanner (1 T field) for 16 s, there can be substantial effects on cognition (van Nierop et al., 2012). This is even though the head motion was about 10 times slower than the molecular motion induced with ultrasound for our strongest stimulus (Figure 3). The peak velocity of the head movements in that study is estimated to be *v* = 0.35 m/s, assuming average head circumference of 0.56m. In *B* = 1T, this yields up to *E* = 0.35 V/m. These effects were noted about 90 s following the head motion, indicating that the Lorentz-based effects can outlive their initiation.

Lstim uniquely combines noninvasiveness, sharp focus, targeting flexibility, and the efficacy of electrical stimulation. No other existing approach has these features. This intersection of desirable features is a result of favorable properties of the two constituting fields. Both magnetic and ultrasonic fields are non-invasive, and the hardware for both is readily available in the form of ultrasonic transducers and MRI scanners, respectively. Lstim owes its sharp focus to focused ultrasound. The spatial resolution of the method is in theory comparable to the voxel size of standard imaging transducer systems. For brain applications, existing hardware is able to focus ultrasound, through the human skull, into deep brain circuits that span only about 3 mm in diameter (Ghanouni et al., 2015). Furthermore, these existing transducer system target ultrasound flexibly, in software. Since ultrasound reaches a target in microseconds, the theoretical temporal resolution of Lstim thus reaches tens-hundreds of microseconds.

Once optimized in regard to stimulation waveforms, this new mode of noninvasive targeted stimulation may develop into a new tool to manipulate neural circuits in the human brain. Such an approach could provide the sorely needed diagnostic tools. For instance, clinical teams could use Lstim to systematically identify the neural circuits that are most strongly involved in specific signs and symptoms in a given patient. The identified circuits could subsequently be modulated using existing clinical tools such as deep brain stimulation or spinal cord stimulation, now applied in a targeted, personalized manner. Analogously, Lstim could be applied in basic neuroscience research as a causal tool to manipulate neural circuits in the human brain. As noted above, ultrasound has been applied to the human brain inside strong magnetic fields (3 T and 7 T) already. These envisioned applications are therefore expected to be safe.

Although we have primarily focused on medical applications, Lstim could be applied to remotely induce electric fields also in other disciplines. For example, the method could be used to remotely catalyze chemical reactions in situations in which a sample cannot be influenced by the insertion of metallic electrodes. Food processing, and remote stimulation of sterile tissue or cell cultures constitute additional examples.

In summary, we asked whether the combination of ultrasonic and magnetic fields could provide a noninvasive alternative to existing deep tissue electrical stimulation methods. This study provides an initial proof of concept using intact human peripheral nervous system. The approach, once optimized for specific kinds of excitable structures, may provide the long-sought tool to manipulate the nervous system in humans noninvasively and in a targeted manner.

## Methods

### Subjects and apparatus

The study was approved by the Institutional Review Board of the University of Utah. Eighteen subjects (6 females, 12 males, aged between 21-38 years) participated in the study. All subjects provided an informed consent.

Subjects were asked to gently rest the thumb of their right hand on a plastic coupling cone filled with degassed water (Figure 2a). The height of the cone was 52 mm and its total diameter 70 mm. The aperture (1 mm-thick plastic) had a 16 mm in diameter opening for the ultrasound beam. A focused, MRI-compatible ultrasonic transducer (H-115, Sonic Concepts, 64 mm diameter, 52 mm focal depth) was positioned 52 mm below the aperture (Figure 2a). The transducer was operated at 258 kHz. Stimuli were generated by a custom Matlab program that produced the stimulation waveforms in a programmable function generator (33520b, Keysight). The signals were amplified using a 55-dB, 258 kHz–30 MHz power amplifier (A150, Electronics & Innovation).

Subjects had their eyes closed and wore noise-cancelling earmuffs (X4A, 3M; 27 dB noise reduction) so that they could fully focus on the stimuli. Subjects could not hear or see the stimuli or their generation.

### Stimuli

The stimulation was performed inside the bore of a 7 T MRI scanner (Bruker BioSpec) or at a 3 m distance away from it. The stimulation order was randomized, without replacement, such that half of the subjects experienced the stimulation in the scanner first and the other half near the scanner first. Subjects were asked to place the finger on the aperture in the direction perpendicular to the ultrasonic and magnetic fields (Figure 2a) to maximize the Lstim effects.

We used 9 distinct stimuli, of 3 pressure levels and 3 distinct waveforms. A tenth, sham stimulus, delivered negligible pressure (5 kPa, corresponding to the noise level of the amplifier-transducer output) under the same conditions. The parameters were chosen to provide safe and effective stimulation. The relatively low carrier frequency (258 kHz) was chosen to maximize the Gildermeister effect (integration of membrane potential toward a threshold), which is more effective at lower frequencies (Ward, 2009). The duration of each stimulus (200 ms) was chosen to provide ample time for the integration. The peak pressure amplitudes of the ultrasound measured at the center of the aperture were 0.35 MPa, 0.53 MPa, and 0.7 MPa. The peak pressures were chosen such as to trigger appreciable electric intensities at target (up to 3.1 V/m), but low enough to comply with the *I*_SPPA_ Track 3 510(k) recommendation for each pulse and within the *I*_SPTA_ recommendation over the course of the experiment (see Stimulus safety), and low enough to prevent unpleasant nociceptive responses. The stimuli were either continuous (200 ms of tone burst) or pulsed at 500 Hz or 10 kHz both 50% duty.

The pressure fields were measured using a capsule hydrophone (HGL-0200, Onda) calibrated between 250 kHz and 40 MHz and secured to 3-degree-of-freedom programmable translation system (Aims III, Onda). There were 10 repetitions of the 10 stimuli, producing a total of 100 stimulation trials per subject inside the scanner and 100 trials outside the scanner. The stimuli were delivered every 8-12 s. The stimuli were drawn from the 100-stimulus set randomly without replacement. This way, stimulus order could not affect the results.

### Responses and their assessment

Subjects were instructed to report a percept with a verbal command of any combination of {Pain, Vibration, Tap}, and their intensity (1: low, 2: medium, 3: high). Following each stimulus, the experimenter was prompted to entered the reported sensation (or lack thereof) and its intensity into a command line of the same Matlab program that scheduled the stimuli. The experimenter was blinded to the stimuli. Following the experiment, for each stimulus type, the response magnitude was computed as the proportion of trials in which subjects’ registered a response, weighted by the reported intensity. Vibration and tap responses were grouped together as tactile.

### Acoustic continuum

The acoustic impedance of water and skin, including soft tissues, are closely matched (1.48 MRayl compared to 1.68 MRayl (Kuhn et al., 2008)). This way, about 99.6% of the energy, 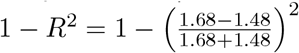, was delivered into the finger. The water-finger interface is therefore essentially acoustically transparent and can be considered as a continuum from the perspective of ultrasound.

### Stimulus safety

The ultrasonic stimuli used in this study were safely below the FDA 510(k) Track 3 recommendations (FDA, 2019). In particular, the highest peak pressure used in the study, 0.7 MPa, corresponds to peak intensity of 15.3 W/cm^2^, which is well below the FDA recommendation of *I*_SPPA_ = 190 W/cm^2^ (Figure 3). In addition, the time-average spatial peak intensity was *I*_SPTA_ = 150 mW/cm^2^, also below the FDA recommendation of *I*_SPTA_ = 720 mW/cm^2^. The computation of the charge density (Figure 3) assumed brain conductivity of 0.26 S/m (Koessler et al., 2017). Thus, stimuli of much higher levels could be used, from both the ultrasound safety and electrical stimulation safety perspectives. The 0.7 MPa maximum allowed all sensations to be well tolerated by the subjects. No subject lifted their finger from the aperture during the experiment. No harmful effects were detected by the experimenter or reported by the subjects.

